# CNVpytor: a tool for CNV/CNA detection and analysis from read depth and allele imbalance in whole genome sequencing

**DOI:** 10.1101/2021.01.27.428472

**Authors:** Milovan Suvakov, Arijit Panda, Colin Diesh, Ian Holmes, Alexej Abyzov

**Affiliations:** Department of Quantitative Health Sciences, Mayo Clinic, Rochester, MN; Center for Individualized Medicine, Mayo Clinic, Rochester, MN; Department of Bioengineering, University of California, Berkeley, USA

**Keywords:** copy number variations, copy number alternations, whole genome sequencing, Python

## Abstract

Detecting copy number variations (CNVs) and copy number alterations (CNAs) based on whole genome sequencing data is important for personalized genomics and treatment. CNVnator is one of the most popular tools for CNV/CNA discovery and analysis based on read depth (RD). Herein, we present an extension of CNVnator developed in Python -- CNVpytor. CNVpytor inherits the reimplemented core engine of its predecessor and extends visualization, modularization, performance, and functionality. Additionally, CNVpytor uses B-allele frequency (BAF) likelihood information from single nucleotide polymorphism and small indels data as additional evidence for CNVs/CNAs and as primary information for copy number neutral losses of heterozygosity. CNVpytor is significantly faster than CNVnator—particularly for parsing alignment files (2 to 20 times faster)—and has (20-50 times) smaller intermediate files. CNV calls can be filtered using several criteria and annotated. Modular architecture allows it to be used in shared and cloud environments such as Google Colab and Jupyter notebook. Data can be exported into JBrowse, while a lightweight plugin version of CNVpytor for JBrowse enables nearly instant and GUI-assisted analysis of CNVs by any user. CNVpytor release and the source code are available on GitHub at https://github.com/abyzovlab/CNVpytor under the MIT license.

## Introduction

The continuous reduction of cost made the whole genome sequencing (WGS) to be widely used in different research projects and clinical applications. Consequently, many approaches for processing, analyzing, and visualizing WGS data have been developed and are being improved. Detection and analysis of copy number variations (CNVs) based on WGS data is one of them. Research directions related to the cancer genomics, single-cell sequencing, and somatic mosaicism create huge amounts of data and demands for processing on the cloud that require further improvements of CNV callers, moving to parallel processing, better compression, modular architecture, and new statistical methods.

CNVnator is a method for CNV analysis based on read depth (RD) of aligned reads. It was determined to have high sensitivity (86%-96%), low false-discovery rate (3%−20%), and high genotyping accuracy (93%-95%) for germline CNVs in a wide range of sizes from a few hundred base pairs to chromosome size events (Abyzov et al., 2011, Mills et al., 2011, Duan et al., 2013, Legault et al., 2015, Trost et al., 2018). Since its development a decade ago, the tool has been widely used in different scientific areas by researchers around the world for detection of CNVs in a variety of species with different genome sizes: bacteria (Coll et al., 2014), fungi (Cabañes et al., 2015), plants (Fuentes et al., 2019, Gordon et al., 2014, Wallace et al., 2014), insects (Choi et al., 2015), fish (Chain et al., 2014), birds (Yi et al., 2014), mammals (Hermsen et al., 2015, Wang et al., 2015, Gokcumen et al., 2013, Pezer et al., 2015) and humans (Abel et al., 2020, Mills et al., 2011, Sudmant et al., 2015, Nagasaki et al., 2015). It has been used to discover somatic variations in cancer and disease studies (Han et al., 2020) and to find mosaic variants in human cells (Guo et al., 2019). Although CNVnator was developed to detect germline CNVs, it is well-suited to discover copy number alteration (CNAs) present in a relatively high (>50%) fraction of cells, such as somatic alteration found in cancers. It was not, however, designed for nor capable of aiding analysis of copy number neutral changes.

Here we describe CNVpytor, a Python extension of CNVnator. CNVpytor inherits the reimplemented core engine of CNVnator and extends visualization, modularization, performance, and functionality. Along with RD data, it enables consideration of allele frequency of single nucleotide polymorphism (SNP) and small indels as an additional source of information for the analysis of CNV/CNA and copy number neutral variations. Along with RD data, this information can be used for genotyping genomic regions and visualization.

## Results

### Analysis of RD signal

CNVpytor inherits the RD analysis approach developed in CNVnator (Abyzov et al., 2011). Briefly, it consists of the following steps: reading alignment file and extracting RD signal, binning RD signal, correcting the signal for GC bias, segmenting the signal using the mean-shift technique, and calling CNVs (**Fig. 1**). RD signal can be parsed from BAM, SAM, or CRAM alignment files and is counted in 100 bp intervals, resulting in a small footprint of intermediate .pytor files in HDF5 format (**Table 1**). Because of using the pysam (Gilman et al., 2019) library for parsing, this step (the most time-consuming one) is parallelized and can be conducted very efficiently, particularly in comparison with the older tool (**Table 1**). The binning step integrates RD over larger bins that are limited to multiples of initially stored 100 bp bins. Next, the bias in the read depth signal due to GC content (so-called GC bias) is adjusted. For the human reference genomes GRCh37 and GRCh38 per bin, GC content is pre-calculated and supplied as a resource with the CNVpytor package. For other genomes, GC content can be calculated during runtime from a provided FASTA file or precalculated and added to the CNVpytor resource for future usage. Once information about read coverage (and variants, see below) is extracted from an alignment file, the following analysis steps take place (i.e., read input and write output) with the same file. As a result, histograms for each processed bin size and information about CNV calls, including coordinates, different statistics, and p-values are all stored in one .pytor file and can be extracted into Excel (TSV file) or a VCF file.

**Figure 1.**
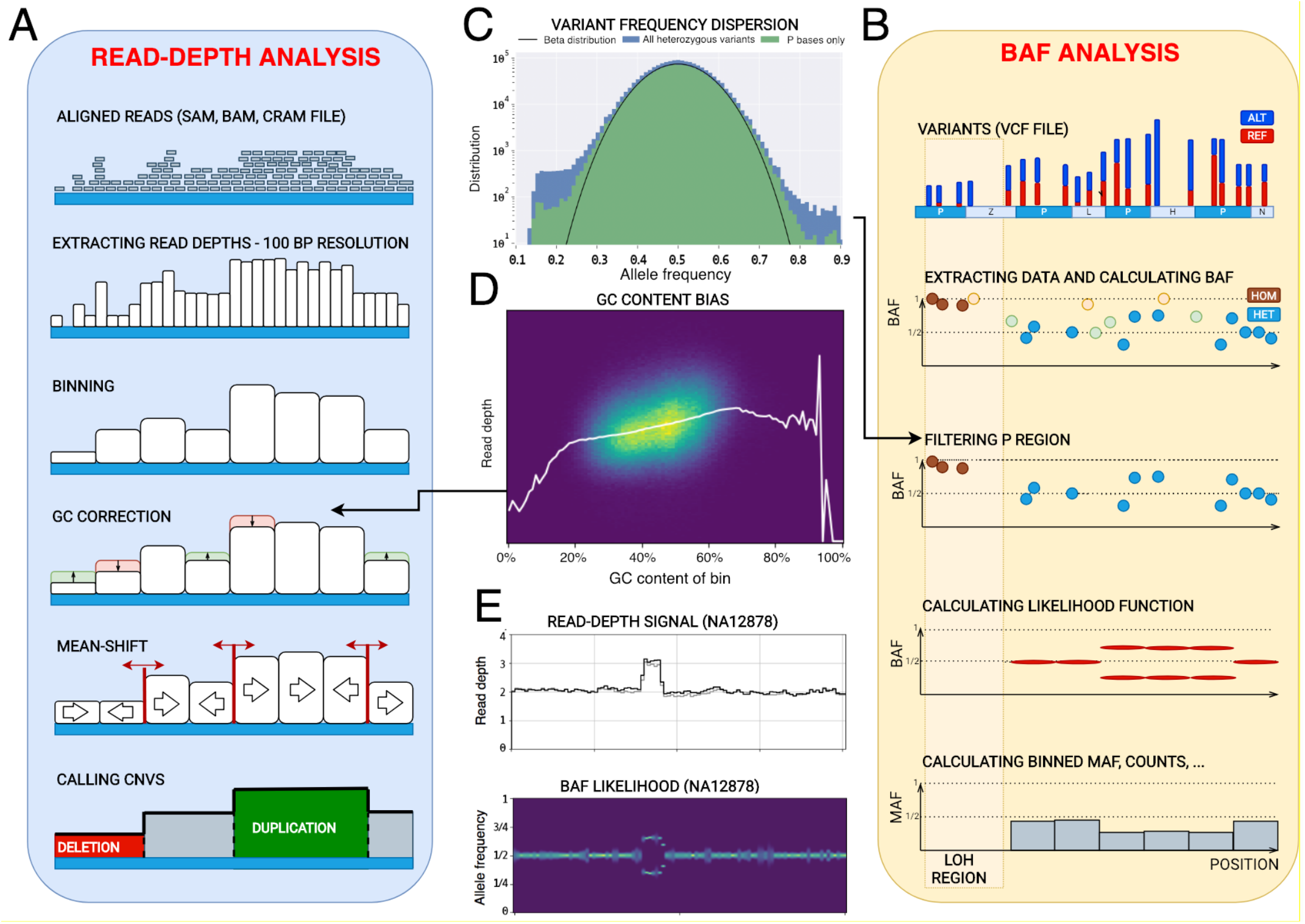
Schematics of core algorithm and data processing steps. (A) Read depth analysis steps include parsing alignment file, calculating and storing read depths in 100 bp intervals, binning using user-specified bin size, correcting RD for GC bias, segmenting by mean-shift, and calling CNVs. (B) B-allele frequency analysis steps include reading variant file storing the data about SNPs and small indels, filtering variants using strict mask, calculating BAF for heterozygous variants (HETs), and calculating likelihood function for bins. For CNVs, BAF signal splits away from value 0.5 expected for HETs. (C) Distribution of the variant allele frequency for all variants and variants within strict mask as defined by the 1000 Genomes Project. Black line shows fit by gaussian distribution. (D) An example of RD depending on GC within bin. Statistics of RD signal within bins of the same percentage of GC content is used to correct for GC bias in the signal. White line represents average RD level for bins with given GC content. (E) An example of RD and BAF signals for a germline duplication in NA12878 sample (raw RD signal is in grey, GC-corrected RD signal is in black, brighter color of BAF likelihood corresponds to higher values of the likelihood).

**Table 1.** Efficiency of parsing alignment file on modern computers in relation to sequencing coverage and engaged number of CPU cores.

| Sequencing coverage of the human genome | Parsing time |  |  |  | File size |  |  |  |
| --- | --- | --- | --- | --- | --- | --- | --- | --- |
|  | CNVnator | CNVpytor |  |  | CNVnator | CNVpytor |  |  |
|  |  | 4 cores | 8 cores | 23 cores |  | RD parsing | BAF parsing | With 1 & 10 kbp bins |
| 5X | 10 min | 5 min | 3 min | <2 min | 1 Gb | 18 Mb | 20 Mb | 250 Mb |
| 30X | 1 h | 28 min | 18 min | 10 min | 1.5 Gb | 19 Mb | 20 Mb | 250 Mb |
| 100X | 3.3 h | 1.5 h | 1 h | 33 min | 2 Gb | 20 Mb | 20 Mb | 250 Mb |

### Analysis of variant data

A novel feature of CNVpytor is the analysis of information from SNPs and small indels. An imbalance in the number of haplotypes can be measured using allele frequencies traditionally referred to as B-allele frequency (BAF) (Peiffer et al., 2006, Loh et al., 2018). The main advantage of using BAF compared to RD is that BAF values do not require normalization and are distributed around 0.5 by binomial distribution for heterozygous variants (HETs). Additionally, BAF is complementary to RD signal, as it changes for copy number neutral events such as loss of heterozygosity. However, BAF dispersion can be measured incorrectly due to systematic misalignment particularly in repeat regions, incomplete reference genome, or site-specific noise in sequencing data. To mitigate this issue, we filtered out HETs in the fraction of genome that is inaccessible to short read technologies, as defined by the strict mask from the 1000 Genomes Project (Auton et al.). Such filtering removes almost all HETs with outlier values of BAF, while values for the retained variants closely follow binomial distribution (**Fig. 1C**). To integrate BAF information within bins, CNVpytor calculates the likelihood function that describes an imbalance between haplotypes (see **Methods**). Currently, BAF information is used when genotyping a specific genomic region where, along with estimated copy number, the output contains the average BAF level and two independent p-values calculated from RD and BAF signal. Variant data can also be plotted in parallel with RD signal (**Fig. 2**). Same as for RD signal, binned information calculated from variants is stored in and can be extracted from .pytor file.

**Figure 2.**
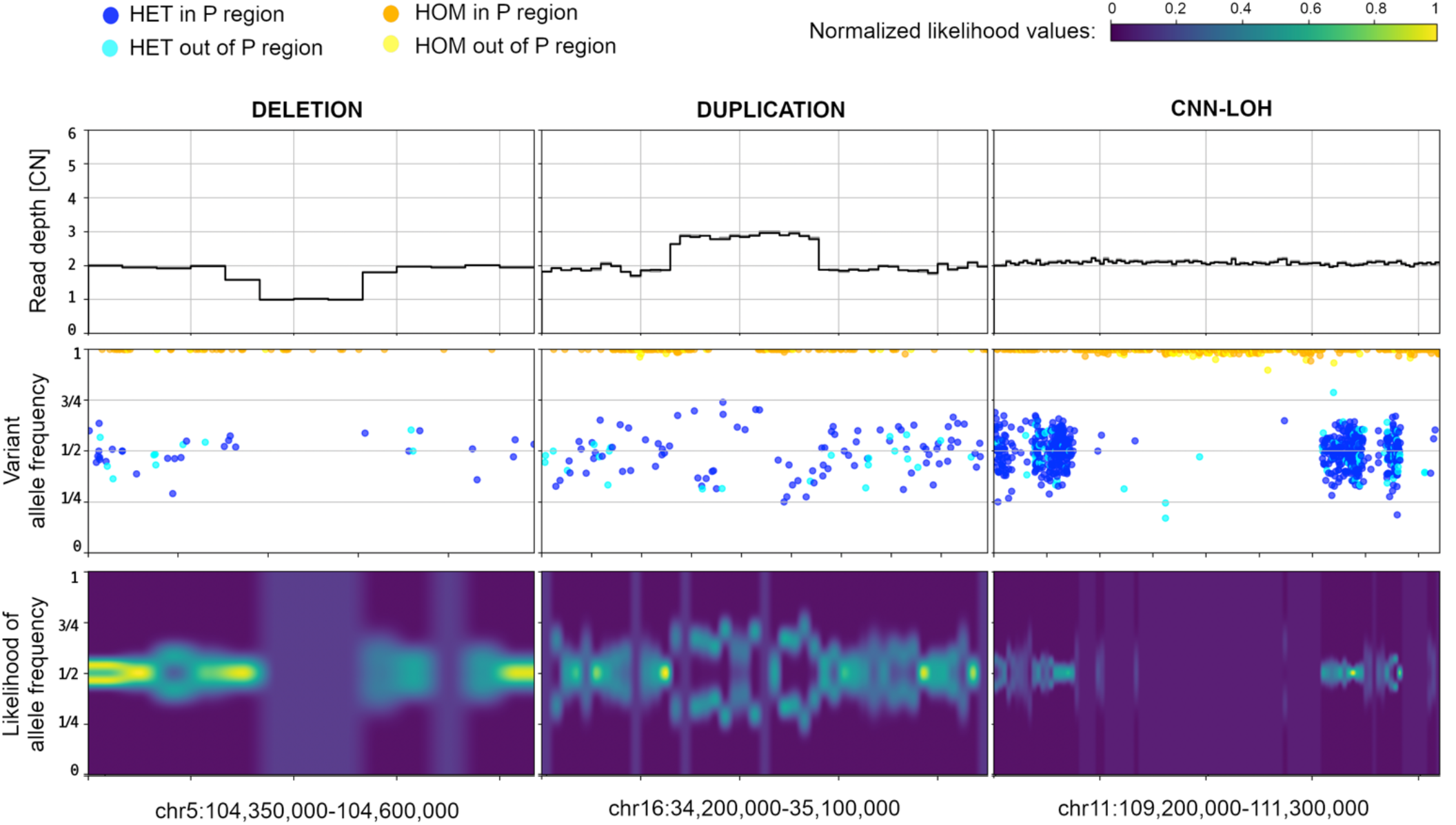
BAF signal corroborates and complements RD signal. Example of CNVpytor *region* plots produced for deletion (left), duplication (middle), and CNN-LOH (right) for NA12878 sample. Within the coordinates of heterozygous deletion, there is a 50% drop in RD signal and a loss in heterozygosity in BAF signal (i.e., no heterozygous SNPs in the region). Duplication of one haplotype results in the increase of RD signal by 50% and in a split in VAF distribution of SNPs and a split in BAF likelihood function. In the CNN-LOH region, few reliable heterozygous SNPs are detected while RD signal does not change. Likelihood function values are normalized to maximal across the range.

### Running CNVpytor

CNVpytor is to be run in a series of steps (**Fig. 3**). For enhanced flexibility, RD and BAF processing workflows proceed in parallel. In this way each workflow can be run at different times or even on different computers. For example, data parsing steps can be run on a cloud where data (i.e., alignment files) is accessible, resulting in less than 25 Mb .pytor files that then can be copied to local computer/cluster where the remaining calculation steps will be performed. If necessary, a user can run additional calculations (e.g., conduct processing with different bin size) using the same .pytor file in the future, allowing for further flexibility in data analysis.

**Figure 3.**
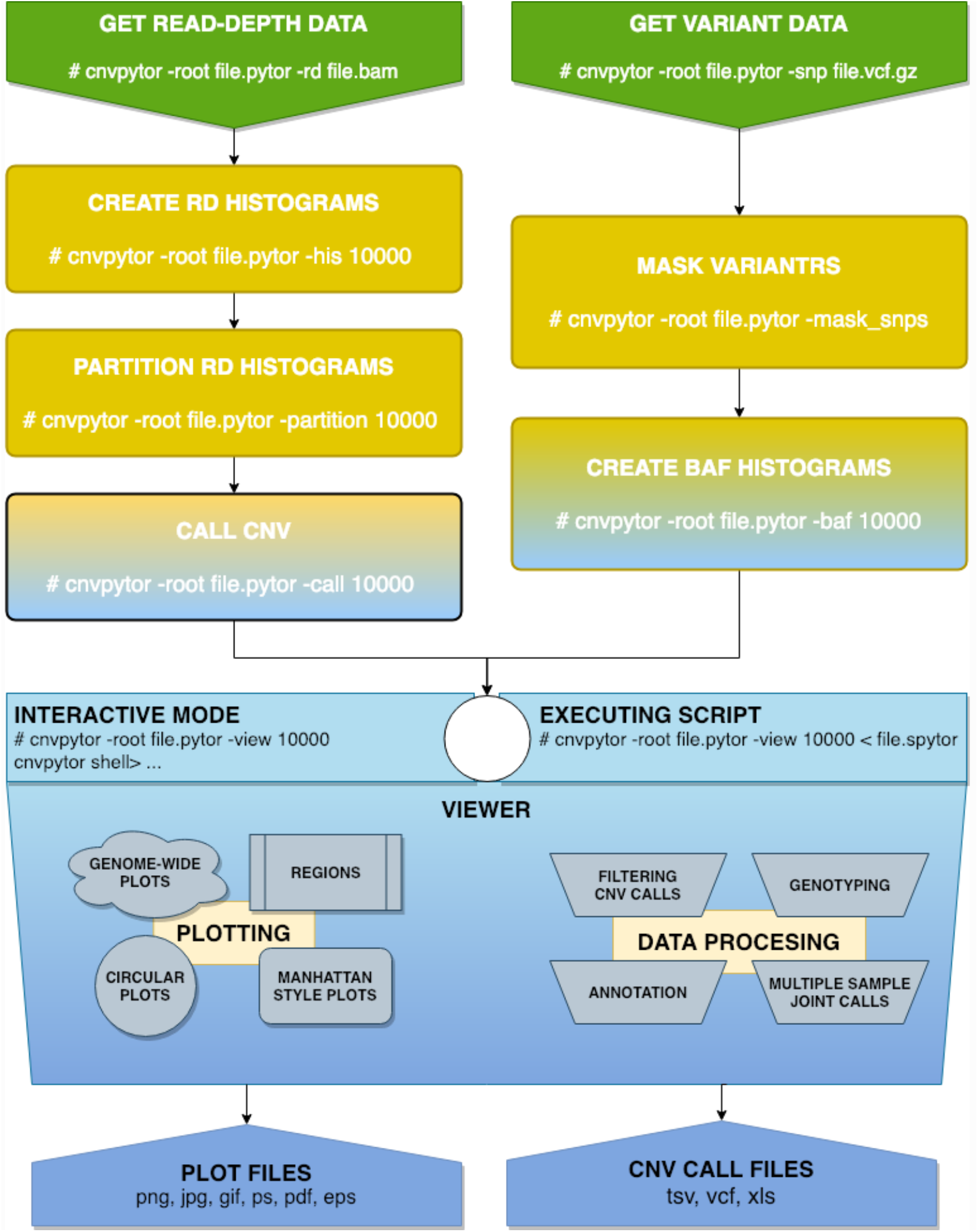
CNVpytor workflow. Steps used in data processing. On the left: reading RD data from alignment file, creating histograms, segmentation, calling CNVs. On the right: reading SNP and indel data from VCF file, filtering variants using strict mask, calculating histograms and likelihood function. Visualization using both RD and/or BAF data can be done from an interactive command line interface or automatically by running script file.

Routine processing steps can be followed by CNV visualization and analysis in the Viewer session, which can be interactive or hands-off. Implementation of interactive mode is inspired by a Linux shell with tab completion and with a help page similar to the man pages. In this mode a user can instantly make various visualizations, preview and filter CNV calls, annotate calls, create joint calls across multiple samples, and genotype specified regions. The viewer does not save results into the .pytor file, and outputs are printed and plotted on the screen or exported to an output file(s). Hand-off mode executes user-written script(s) with CNVpytor commands. Such scripts can be used as part of the processing pipeline where, for example, images of signals around called CNVs are generated and stored for possible future inspection. Through the viewer interface, it is possible to directly access Python and run code. This allows user to access some standard features of underlying libraries, e.g., matplotlib library can be used to customize plots.

CNVpytor can be used as the Python module. All functionalities, like reading and editing CNVpytor data files, and all calculation steps and visualizations can be performed by calling functions or classes. This way CNVpytor can be easily integrated in different platforms and computing environments; for example, CNVpytor can be run from Jupyter Notebook on a local machine or in cloud services, e.g., Google Colab.

### Visualizations

Visualization of multiple tracks/signals can be done interactively by mouse and by typing relevant commands as well as by running scripts with CNVpytor commands provided as inputs to CNVpytor. CNVpytor has extended visualization capabilities with multiple novel features as compared to CNVnator. For each sample (i.e., input file) multiple data tracks such as RD signal, BAF of SNPs, and binned BAF likelihood can be displayed in an adjustable grid layout as specified by a user. Specifically, multiple regions across multiple samples can be plotted in parallel, facilitating comparison across samples and different genomic loci (**Fig. 2**). To get a global view, a user can visualize an entire genome in a linear or circular fashion (**Fig. 4, S1, S2**). Such a view can be useful in judging the quality of samples and in visually checking for aneuploidies.

**Figure 2.**
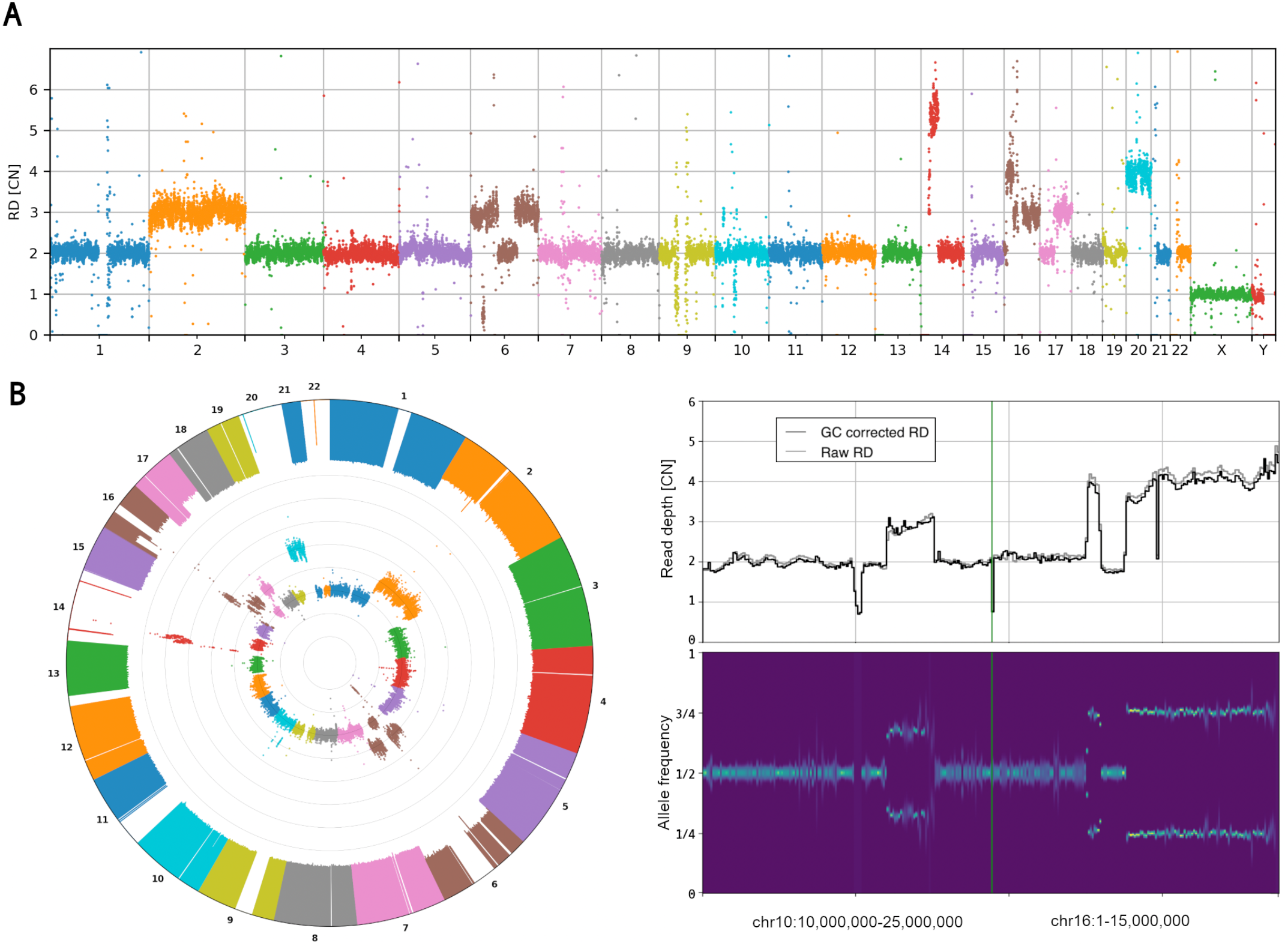
Novel features of visualization using data for HepG2 cell line. Genome-wide visualizations can be useful in some cases, like cancer studies or clinical applications like screening for chromosomal abnormalities. They also can be useful for quick insight in sequencing or single-cell amplification quality. To demonstrate genome-wide plot types, HapG2 immortal cell line sample is used. A) Manhattan and B) circular style plot of RD signal. Large CNVs and chromosome copy number changes are apparent. One can also judge about dispersion of the RD signal. The inner circle in B shows RD signal while outer shows minor allele frequency (MAF). Regions where we can see loss of MAF signal with normal RD, e.g., chromosome 14 or 22, are chromosomal CNN-LOH. C) Examples of smaller CNVs not apparent in the global view. Each CNV is about 15 Mbp. The displayed regions with multiple CNVs (that are possibly complex events) were hardly visible on the circular plot.

Some additional features include GC-bias curve plot and 2D histogram (**Fig. 1D**), allele frequency distribution per region (**Fig. 1C**), and comparing RD distributions between two regions (**Fig. S3**). Figure resolution, layout grid, colors, marker size, titles, and plotting style are adjustable by the user.

### Integration with JBrowse

CNVpytor also has implemented functionality to export data into formats that can be embedded into JBrowse, a web-based genome browser used to visualize multiple related data tracks (**Fig. S4**). The export enables users to utilize JBrowse capabilities to visualize, compare, and cross-reference CNV calls with other data types (such as CHiP-seq, RNA-seq, ATAC-seq, etc.) and annotations across genome. Exported data provide 3 resolutions of RD and BAF tracks (1 kbp, 10 kbp, and 100 kbp bins) while the appropriate resolution is chosen automatically by JBrowse depending on the size of the visualized genomic region. Multiple .pytor files can be exported at once.

Alternatively, a user can utilize a lightweight CNVpytor plugin for JBrowse. The plugin takes information about coverage from a relatively small (as compared to BAM) VCF file and on the fly performs the read depth and BAF estimation, segmentation, and calling. For read depth analysis, the plugin fetches the information from the DP field in the VCF field and uses it as a proxy for actual coverage. Since for large bin sizes such an estimate corresponds well to the actual value (**Fig. S5**), the plugin enables quick and easy review of large copy number changes in a genome. For BAF analysis, the plugin conducts analyses the same way as a stand-alone application. All temporary values are stored in the browser cache for fast and interactive visualization of a genomic segment. As well as improving responsiveness by eliminating the network lag of a client-server application, this ensures that no information about a personal genome is transferred to external servers. Once the analyses are complete, the results are instantly visualized using JBrowse’s native capabilities. Usage cases of the plugin are: 1) quick and visual cross-referencing of copy number profiles between multiple samples and in relation to other data types, and 2) a review of a personal genome(s) for large CNVs in a simple user-friendly environment.

## Conclusion

Development of new and maintenance and improvement of existing bioinformatics tools are driven by changing data types, demands for newer and user-verifiable analyses, necessity for processing larger datasets, and the evolving nature of computational infrastructures and platforms. CNVpytor brings the functionality of its predecessor CNVnator to a new level and significantly expands it. CNVpytor is faster and virtually effortless to install, requires minimum space for storage, enables analysis of BAF, provides users with extended visualization and convenient functionality for result curation, and is equipped for integration with other tools. More than that, CNVpytor establishes a framework for discovering and analyzing copy number changes from whole genome sequencing data either by an individual researcher or clinician or in a collaborative and shared environment.

## Materials and Methods

### RD analysis

Calculations for RD binning, mean-shift algorithm, partitioning and calling CNVs are explained in details in CNVnator paper (Abyzov et al., 2011). The only difference in CNVpytor implementation is how information about GC content is obtained. For two versions GRCh37 (hg19) and GRCh38 of the human reference genome information about GC and AT content for each 100 base pair bins is provided as resource data within CNVpytor package. This way user does not need to have reference genome FASTA for GC correction.

### Variant data

CNVpytor imports information about SNPs and single letter indels form variant (VCF) file. All other variants are ignored. For each variant following data is stored in CNVpytor file: chromosome, position, reference base, alternative base, reference count (*ref_i_*), alternative count (*alt_i_*), quality, and genotype (0/1 or 1/1).

In normal case with one copy of each haplotype, heterozygous SNPs are visible in half of the reads coming from that haplotype. In other words, B-allele frequency distributes around 1/2. Contrarily, in the regions with constitutional duplication of one haplotype, heterozygous SNPs are expected with frequency equals 2/3 or 1/3 depending are they located on duplicated haplotype or not. This split from value 1/2 can be visible in plot of BAFs vs position of variants as shown in the right panel of Figure 1. Similarly, for homozygous deletion complete loss of heterozygous SNPs is expected. In the case of somatic sub clonal CNA (e.g., frequently observed in cancer genomes), the ratio between haplotypes can be an arbitrary number depending on cell frequencies with the CNA. Consequently, the split in BAF plot varies from 0 through 1. Measuring the level of this split can provide useful information about type of CNV. Moreover, copy number neutral loss of heterozygosity (CNN-LOH) can be detected this way.

For each stored variant we can calculate two frequencies defined in following way:

- B-allele frequency (BAF): 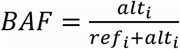
- Minor allele frequency: 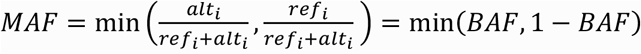

One of next generation sequencing features is that some bases are not accessible for variant discovery using short reads, due to repetitive nature of the human genome. In the 1000 Genomes Project, genome mask is created to tabulate bases for variant discovery. There are about 74% of bases marked passed (P), what correspond to about 77% of non-N bases (1000 Genomes Project Consortium, 2015). We use that mask to filter out variants called in non-P regions. This way we eliminate around 22% of variants but there is benefit because with less false-positive heterozygous SNPs statistics is improved (**Fig. 1C**) and this improves quality of further calculations. The same way as GC content, information about strict mask P regions for two versions of human reference genome are stored in resource files that are part of CNVpytor package.

### Calculating BAF likelihood function

The ratio between reads coming from one or another haplotype is distributed following binomial distribution. If counts are known then one can calculate likelihood function for that ratio:

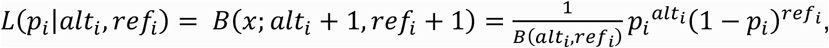

where *p_i_* is allele frequency for variant *i* and *B*(*alt_i_, ref_i_*) is normalization constant.

By multiplying likelihood functions of individual variants in a bin one can obtain likelihood for each bin. This is true only if real value of that ratio does not change within a that bin. However, we are using non-phased variant counts, what means that there is 50% of chance that variant is coming from one or another haplotype. Frequency of a fixed haplotype is sometimes described by either BAF and 1 - BAF distributions.

In that case we have to use symmetrized beta function for likelihood:

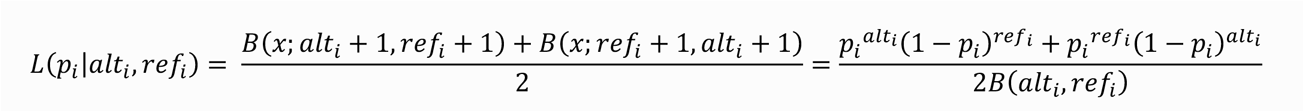

Likelihood function for each bin is calculated as a product of individual likelihood functions of variants within that bin:

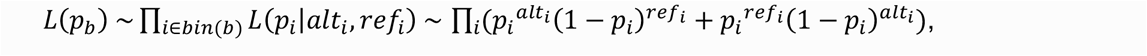

where *p_b_* is allele frequency for bin *b*. To calculate likelihood functions we use discretization. Interval [0,0.5] is discretized using some resolution (default is 101 point) and for each point function is calculated by multiplying values of symmetrized beta functions for each variant. Position of maximum likelihood represents most probable BAF value in particular bin. Along with likelihood function average values of variant BAF and MAF are calculated per bin and stored in CNVpytor file together with counts of homozygous and heterozygous variants.

### Filtering CNV calls

For each CNV call following values are calculated: (1) event type: “deletion” or “duplication”; (2) coordinates in the reference genome; (3) CNV size; (4) RD normalized to 1; (5) e-val1 – p value calculated using t-test statistics between RD difference in the region and global (i.e. across whole genome) mean; (6) e-val2 – p value from the probability of RD values within the region to be in the tails of a gaussian distribution of binned RD; (7) e-val3 – same as e-val1 but without first and last bin; (8) e-val4 – same as e-val2 but without first and last bin; (9) q0 – fraction of reads mapped with zero quality within call region; (10) pN – fraction of N bases (i.e., unassembled reference genome) within call region; (11) dG – distance to nearest gap in reference genome.

There are five parameters in viewer mode used for filtering calls: CNV size, e-val1, q0, pN and dG. Those parameters will define which calls CNVpytor will plot or print out. When calls are printed or exported CNVpytor optionally can generate graphical file(s) with plot of CNV call region containing user specified tracks.

### Annotating CNV calls

To annotate called regions we use Ensembl REST API (overlap/region resource). It is an optional step requires web connection and is executed when calls are previewed by user or exported to an output file. The annotation is added in an additional column in the output and contains string with gene names, Ensembl gene IDs and with information about position of genes relative to CNV (i.e., inside, covering, or intersecting left/right breakpoints of CNV region).

### Genotyping

Copy number of a provided genomic region is calculated as a mean RD within the region divided by mean autosomal RD scaled by 2. To achieve better precession, first and last bin content are weighted by the fraction of overlap with the provided region. Optionally CNVpytor can provide additional values: (1) e-value from the probability of RD values within the region to be in the tails of a gaussian distribution of binned RD (analogous to e-val2); (2) q0 – fraction of reads mapped with q0 quality within call region; (3) pN – fraction of reference genome gaps (Ns) within call region; (4) BAF level estimated using maximum likelihood method; (5) number of homozygous variants within the region; (6) number of heterozygous variants within the region; (7) p-value based on BAF signal.

### Merging calls over multiple regions

To make joint call set for multiple samples CNVpytor procceds in the following way:

1. Filter calls using user defined ranges for size, p-val, q0, pN and dG;
2. Sort all calls for all samples by start coordinate;
3. Select first call in that list that is not already processed and select calls from other samples with the reciprocal overlap larger than 50%;
4. For selected calls, calculate genotypes within the region of intersection and, optionally, annotate with overlapping genes.

If specified, for each joint CNV call CNVpytor will create graphical file with plot of call region containing user specified tracks.

### Data format and compression

For data storage and compression, we used HDF5 file format and h5py python library. Additional compression is obtained by storing RD signal using 100 base pair bins. The same bin size is used for storing reference genome AT, GC and N content. Data organization within .pytor file is implemented in IO module which can be used to open and read different datasets from an external application. Python library xlwt is used to generate spreadsheet files compatible with Microsoft Excel.

### Visualizations

Matplotlib (Hunter, 2007) python library is used for creating and storing visualizations. Different plotting styles available within matplotlib. Derived installed libraries can be used as well as variety of file formats for storing graphical data.

### Module dependences

CNVpytor depends on several widely used python packages: requests 2.0 or higher, gnureadline, pathlib 1.0 or higher, pysam 0.15 or higher, numpy 1.16 or higher, scipy 1.1 or higher, matplotlib 2.2 or higher, h5py 2.9 or higher, xlwt 1.3 or higher. All dependences are available through pip installer what makes installation of CNVpytor straightforward.

### Software availability

Straightforward installation with Pip, Conda or Docker tool makes CNVpytor available on different operating systems. As python module, CNVpytor is ready to embed in different platforms and computing environments. CNVpytor source code is available on the GitHub (https://github.com/abyzovlab/CNVpytor) and can be installed via pip tool (https://pypi.org/project/CNVpytor), the BioConda Project or the Docker. It is released under MIT license. The user guide with API documentation is available on the project’s GitHub page.

## Acknowledgement

We thank Abyzov lab members for useful discussions and suggestions. We are grateful to users on the GitHub for useful suggestions and bug reports.

## Notes

### Competing Interest Statement

The authors have declared no competing interest.

## References

Abel, H. J., Larson, D. E., Regier, A. A., Chiang, C., Das, I., Kanchi, K. L., Layer, R. M., Neale, B. M., Salerno, W. J., Reeves, C., Buyske, S., Matise, T. C., Muzny, D. M., Zody, M. C., Lander, E. S., Dutcher, S. K., Stitziel, N. O. & Hall, I. M. 2020. Mapping and characterization of structural variation in 17,795 human genomes. Nature, 583, 83–89.

Abyzov, A., Mariani, J., Palejev, D., Zhang, Y., Haney, M. S., Tomasini, L., Ferrandino, A. F., Rosenberg Belmaker, L. A., Szekely, A., Wilson, M., Kocabas, A., Calixto, N. E., Grigorenko, E. L., Huttner, A., Chawarska, K., Weissman, S., Urban, A. E., Gerstein, M. & Vaccarino, F. M. 2012. Somatic copy number mosaicism in human skin revealed by induced pluripotent stem cells. Nature, 492, 438–42.

Abyzov, A., Urban, A. E., Snyder, M. & Gerstein, M. 2011. CNVnator: an approach to discover, genotype, and characterize typical and atypical CNVs from family and population genome sequencing. Genome Res, 21, 974–84.

Auton, A., Brooks, L. D., Durbin, R. M., Garrison, E. P., Kang, H. M., Korbel, J. O., Marchini, J. L., Mccarthy, S., Mcvean, G. A. & Abecasis, G. R. 2015. A global reference for human genetic variation. Nature, 526, 6874.

CabañEs, F. J., Sanseverino, W., CastellÁ, G., Bragulat, M. R., Cigliano, R. A. & SÁNchez, A. 2015. Rapid genome resequencing of an atoxigenic strain of Aspergillus carbonarius. Sci Rep, 5, 9086.

Chain, F. J., Feulner, P. G., Panchal, M., Eizaguirre, C., Samonte, I. E., Kalbe, M., Lenz, T. L., Stoll, M., Bornberg-Bauer, E., Milinski, M. & Reusch, T. B. 2014. Extensive copy-number variation of young genes across stickleback populations. PLoS Genet, 10, e1004830.

Choi, J. Y., Bubnell, J. E. & Aquadro, C. F. 2015. Population Genomics of Infectious and Integrated Wolbachia pipientis Genomes in Drosophila ananassae. Genome Biol Evol, 7, 2362–82.

Coll, F., Preston, M., Guerra-AssunÇÃO, J. A., Hill-Cawthorn, G., Harris, D., PerdigÃO, J., Viveiros, M., Portugal, I., Drobniewski, F., Gagneux, S., Glynn, J. R., Pain, A., Parkhill, J., Mcnerney, R., Martin, N. & Clark, T. G. 2014. PolyTB: a genomic variation map for Mycobacterium tuberculosis. Tuberculosis (Edinb), 94, 346–54.

Duan, J., Zhang, J. G., Deng, H. W. & Wang, Y. P. 2013. Comparative studies of copy number variation detection methods for next-generation sequencing technologies. PLoS One, 8, e59128.

Fuentes, R. R., Chebotarov, D., Duitama, J., Smith, S., De La Hoz, J. F., Mohiyuddin, M., Wing, R. A., Mcnally, K. L., Tatarinova, T., Grigoriev, A., Mauleon, R. & Alexandrov, N. 2019. Structural variants in 3000 rice genomes. Genome Res, 29, 870–880.

Gilman, P., Janzou, S., Guittet, D., Freeman, J., Diorio, N., Blair, N., Boyd, M., Neises, T. & Wagner, M. 2019. PySAM (Python Wrapper for System Advisor Model” SAM”). National Renewable Energy Lab.(NREL), Golden, CO (United States).

Gokcumen, O., Tischler, V., Tica, J., Zhu, Q., Iskow, R. C., Lee, E., Fritz, M. H., Langdon, A., StÜTz, A. M., Pavlidis, P., Benes, V., Mills, R. E., Park, P. J., Lee, C. & Korbel, J. O. 2013. Primate genome architecture influences structural variation mechanisms and functional consequences. Proc Natl Acad Sci U S A, 110, 15764–9.

Goldhammer, N., Kim, J., Timmermans-Wielenga, V. & Petersen, O. W. 2019. Characterization of organoid cultured human breast cancer. Breast Cancer Res, 21, 141.

Gordon, S. P., Priest, H., Des Marais, D. L., Schackwitz, W., Figueroa, M., Martin, J., Bragg, J. N., Tyler, L., Lee, C. R., Bryant, D., Wang, W., Messing, J., Manzaneda, A. J., Barry, K., Garvin, D. F., Budak, H., Tuna, M., Mitchell-Olds, T., Pfender, W. F., Juenger, T. E., Mockler, T. C. & Vogel, J. P. 2014. Genome diversity in Brachypodium distachyon: deep sequencing of highly diverse inbred lines. Plant J, 79, 361–74.

Guo, H., Duyzend, M. H., Coe, B. P., Baker, C., Hoekzema, K., Gerdts, J., Turner, T. N., Zody, M. C., Beighley, J. S., Murali, S. C., Nelson, B. J., Bamshad, M. J., Nickerson, D. A., Bernier, R. A. & Eichler, E. E. 2019. Genome sequencing identifies multiple deleterious variants in autism patients with more severe phenotypes. Genet Med, 21, 1611–1620.

Han, L., Zhao, X., Benton, M. L., Perumal, T., Collins, R. L., Hoffman, G. E., Johnson, J. S., Sloofman, L., Wang, H. Z., Stone, M. R., Brennand, K. J., Brand, H., Sieberts, S. K., Marenco, S., Peters, M. A., Lipska, B. K., Roussos, P., Capra, J. A., Talkowski, M. & Ruderfer, D. M. 2020. Functional annotation of rare structural variation in the human brain. Nat Commun, 11, 2990.

Hermsen, R., De Ligt, J., Spee, W., Blokzijl, F., Schôfer, S., Adami, E., Boymans, S., Flink, S., Van Boxtel, R., Van Der Weide, R. H., Aitman, T., Hübner, N., Simonis, M., Tabakoff, B., Guryev, V. & Cuppen, E. 2015. Genomic landscape of rat strain and substrain variation. BMC Genomics, 16, 357.

Hunter, J. D. 2007. Matplotlib: A 2D graphics environment. Computing in science & engineering, 9, 90–95.

Legault, M. A., Girard, S., Lemieux Perreault, L. P., Rouleau, G. A. & Dubé, M. P. 2015. Comparison of sequencing based CNV discovery methods using monozygotic twin quartets. PLoS One, 10, e0122287.

Loh, P. R., Genovese, G., Handsaker, R. E., Finucane, H. K., Reshef, Y. A., Palamara, P. F., Birmann, B. M., Talkowski, M. E., Bakhoum, S. F., Mccarroll, S. A. & Price, A. L. 2018. Insights into clonal haematopoiesis from 8,342 mosaic chromosomal alterations. Nature, 559, 350–355.

Mills, R. E., Walter, K., Stewart, C., Handsaker, R. E., Chen, K., Alkan, C., Abyzov, A., Yoon, S. C., Ye, K., Cheetham, R. K., Chinwalla, A., Conrad, D. F., Fu, Y., Grubert, F., Hajirasouliha, I., Hormozdiari, F., Iakoucheva, L. M., Iqbal, Z., Kang, S., Kidd, J. M., Konkel, M. K., Korn, J., Khurana, E., Kural, D., Lam, H. Y., Leng, J., Li, R., Li, Y., Lin, C. Y., Luo, R., Mu, X. J., Nemesh, J., Peckham, H. E., Rausch, T., Scally, A., Shi, X., Stromberg, M. P., StÜTz, A. M., Urban, A. E., Walker, J. A., Wu, J., Zhang, Y., Zhang, Z. D., Batzer, M. A., Ding, L., Marth, G. T., Mcvean, G., Sebat, J., Snyder, M., Wang, J., Ye, K., Eichler, E. E., Gerstein, M. B., Hurles, M. E., Lee, C., Mccarroll, S. A. & Korbel, J. O. 2011. Mapping copy number variation by population-scale genome sequencing. Nature, 470, 59–65.

Nagasaki, M., Yasuda, J., Katsuoka, F., Nariai, N., Kojima, K., Kawai, Y., Yamaguchi-Kabata, Y., Yokozawa, J., Danjoh, I., Saito, S., Sato, Y., Mimori, T., Tsuda, K., Saito, R., Pan, X., Nishikawa, S., Ito, S., Kuroki, Y., Tanabe, O., Fuse, N., Kuriyama, S., Kiyomoto, H., Hozawa, A., Minegishi, N., Douglas Engel, J., Kinoshita, K., Kure, S., Yaegashi, N. & Yamamoto, M. 2015. Rare variant discovery by deep wholegenome sequencing of 1,070 Japanese individuals. Nat Commun, 6, 8018.

Peiffer, D. A., Le, J. M., Steemers, F. J., Chang, W., Jenniges, T., Garcia, F., Haden, K., Li, J., Shaw, C. A., Belmont, J., Cheung, S. W., Shen, R. M., Barker, D. L. & Gunderson, K. L. 2006. High-resolution genomic profiling of chromosomal aberrations using Infinium whole-genome genotyping. Genome Res, 16, 1136–48.

Pezer, Ž., Harr, B., Teschke, M., Babiker, H. & Tautz, D. 2015. Divergence patterns of genic copy number variation in natural populations of the house mouse (Mus musculus domesticus) reveal three conserved genes with major population-specific expansions. Genome Res, 25, 1114–24.

Sekar, S., Tomasini, L., Proukakis, C., Bae, T., Manlove, L., Jang, Y., Scuderi, S., Zhou, B., Kalyva, M., Amiri, A., Mariani, J., Sedlazeck, F. J., Urban, A. E., Vaccarino, F. M. & Abyzov, A. 2020. Complex mosaic structural variations in human fetal brains. Genome Res, 30, 1695–1704.

Sudmant, P. H., Rausch, T., Gardner, E. J., Handsaker, R. E., Abyzov, A., Huddleston, J., Zhang, Y., Ye, K., Jun, G., Fritz, M. H., Konkel, M. K., Malhotra, A., StÜTz, A. M., Shi, X., Casale, F. P., Chen, J., Hormozdiari, F., Dayama, G., Chen, K., Malig, M., Chaisson, M. J. P., Walter, K., Meiers, S., Kashin, S., Garrison, E., Auton, A., Lam, H. Y. K., Mu, X. J., Alkan, C., Antaki, D., Bae, T., Cerveira, E., Chines, P., Chong, Z., Clarke, L., Dal, E., Ding, L., Emery, S., Fan, X., Gujral, M., Kahveci, F., Kidd, J. M., Kong, Y., Lameijer, E. W., Mccarthy, S., Flicek, P., Gibbs, R. A., Marth, G., Mason, C. E., Menelaou, A., Muzny, D. M., Nelson, B. J., Noor, A., Parrish, N. F., Pendleton, M., Quitadamo, A., Raeder, B., Schadt, E. E., Romanovitch, M., Schlattl, A., Sebra, R., Shabalin, A. A., Untergasser, A., Walker, J. A., Wang, M., Yu, F., Zhang, C., Zhang, J., Zheng-Bradley, X., Zhou, W., Zichner, T., Sebat, J., Batzer, M. A., Mccarroll, S. A., Mills, R. E., Gerstein, M. B., Bashir, A., Stegle, O., Devine, S. E., Lee, C., Eichler, E. E. & Korbel, J. O. 2015. An integrated map of structural variation in 2,504 human genomes. Nature, 526, 75–81.

Trost, B., Walker, S., Wang, Z., Thiruvahindrapuram, B., Macdonald, J. R., Sung, W. W. L., Pereira, S. L., Whitney, J., Chan, A. J. S., Pellecchia, G., Reuter, M. S., Lok, S., Yuen, R. K. C., Marshall, C. R., Merico, D. & Scherer, S. W. 2018. A Comprehensive Workflow for Read Depth-Based Identification of Copy-Number Variation from Whole-Genome Sequence Data. Am J Hum Genet, 102, 142–155.

Wallace, J. G., Bradbury, P. J., Zhang, N., Gibon, Y., Stitt, M. & Buckler, E. S. 2014. Association mapping across numerous traits reveals patterns of functional variation in maize. PLoS Genet, 10, e1004845.

Wang, H., Wang, C., Yang, K., Liu, J., Zhang, Y., Wang, Y., Xu, X., Michal, J. J., Jiang, Z. & Liu, B. 2015. Genome Wide Distributions and Functional Characterization of Copy Number Variations between Chinese and Western Pigs. PLoS One, 10, e0131522.

Yi, G., Qu, L., Liu, J., Yan, Y., Xu, G. & Yang, N. 2014. Genome-wide patterns of copy number variation in the diversified chicken genomes using next-generation sequencing. BMC Genomics, 15, 962.

